# Ablation of *Sam50* is associated with fragmentation and alterations in metabolism in murine and human myotubes

**DOI:** 10.1101/2023.05.20.541602

**Authors:** Bryanna Shao, Mason Killion, Ashton Oliver, Chia Vang, Faben Zeleke, Kit Neikirk, Zer Vue, Edgar Garza-Lopez, Jian-qiang Shao, Margaret Mungai, Jacob Lam, Qiana Williams, Christopher T. Altamura, Aaron Whiteside, Kinuthia Kabugi, Jessica McKenzie, Alice Koh, Estevão Scudese, Larry Vang, Andrea G. Marshall, Amber Crabtree, Janelle I. Tanghal, Dominique Stephens, Ho-Jin Koh, Brenita C. Jenkins, Sandra A. Murray, Anthonya T. Cooper, Clintoria Williams, Steven M. Damo, Melanie R. McReynolds, Jennifer A. Gaddy, Celestine N. Wanjalla, Heather K. Beasley, Antentor Hinton

## Abstract

The Sorting and Assembly Machinery (SAM) Complex is responsible for assembling β-barrel proteins in the mitochondrial membrane. Comprising three subunits, Sam35, Sam37, and Sam50, the SAM complex connects the inner and outer mitochondrial membranes by interacting with the mitochondrial contact site and cristae organizing system (MICOS) complex. Sam50, in particular, stabilizes the mitochondrial intermembrane space bridging (MIB) complex, which is crucial for protein transport, respiratory chain complex assembly, and regulation of cristae integrity. While the role of Sam50 in mitochondrial structure and metabolism in skeletal muscle remains unclear, this study aims to investigate its impact. Serial block-face-scanning electron microscopy (SBF-SEM) and computer-assisted 3D renderings were employed to compare mitochondrial structure and networking in *Sam50*-deficient myotubes from mice and humans with wild-type (WT) myotubes. Furthermore, autophagosome 3D structure was assessed in human myotubes. Mitochondrial metabolic phenotypes were assessed using Gas Chromatography-Mass Spectrometry-based metabolomics to explore differential changes in WT and *Sam50*-deficient myotubes. The results revealed increased mitochondrial fragmentation and autophagosome formation in *Sam50*-deficient myotubes compared to controls. Metabolomic analysis indicated elevated metabolism of propanoate and several amino acids, including ß-Alanine, phenylalanine, and tyrosine, along with increased amino acid and fatty acid metabolism in *Sam50*-deficient myotubes. Furthermore, impairment of oxidative capacity was observed upon *Sam50* ablation in both murine and human myotubes, as measured with the XF24 Seahorse Analyzer. Collectively, these findings support the critical role of Sam50 in establishing and maintaining mitochondrial integrity, cristae structure, and mitochondrial metabolism. By elucidating the impact of *Sam50*-deficiency, this study enhances our understanding of mitochondrial function in skeletal muscle.

## Introduction

Mitochondria are generally associated with their bioenergetic roles in the cell, as they are the hubs of oxidative phosphorylation (1,2). Aside from energy production, mitochondria are also integral in other processes such as maintaining calcium homeostasis and apoptosis (2,3). Being such a functionally important organelle, mitochondria are also implicated in the pathophysiology of numerous diseases when they become dysfunctional. Understanding the processes involved in mitochondrial dysfunction may provide novel therapeutic treatment options for many common diseases. For example, Diabetes mellitus is considered a mitochondrial disease (4) and increasing the understanding of mitochondrial function and morphology in normal and pathophysiological functions may be key in developing novel protocols to treat and even prevent disease. This is exemplified when considering reactive oxidative species (ROS) signaling which may alter various mitochondrial functions including ATP production, which may be linked to the development of metabolic diseases and structural changes (5). Mutations or changes in the function of the mitochondrial DNA (mtDNA) can result in altered mitochondrial function; this can have consequences on insulin production, lead to hypertension, and ultimately be related to the progression of Diabetes Mellitus (6,7). Thus, the prospect of understanding how changes in mitochondrial structure may affect metabolism and remains pertinent for discovering new therapeutic treatments for diseases with high morbidity and mortality (8).

Critical to the functioning of mitochondria are the structural folds of the inner membrane known as cristae (9). Cristae provide a means for mitochondria to maximize the amount of oxidative phosphorylation machinery that can be present relative to the internal volume (9). A key determining factor in mitochondrial ultrastructure is mitochondrial dynamics which consist of continuous cycles of fusion and fission. Mitochondrial dynamics give rise to various mitochondrial phenotypes which range from the typical spherical to stress-states including megamitochondria and donut-shaped (10,11). GTPases facilitate the dynamic events with Mitofusin-1 (Mfn1) and Mitofusin-2 (Mfn2) fusing the outer membrane and Optic atrophy 1 (Opa1) fusing the inner membrane (9,12,13). Opa1 has been shown to be linked to mitochondrial fragmentation while also affecting cristae dimensions, shapes, and size (14). Specifically, Opa1 exists in two principal isoforms, the short and long forms, which believed to be responsible for prompting fission and the fusion of the inner membrane, respectively (15). The maintenance of the proper long vs short isoforms are crucial for mitochondrial health and proper cell signaling (16). Thus, understanding the mechanisms of how modulators of cristae affect overall mitochondrial structure remains important and is still not completely understood.

Mitochondria can be separated into several distinct compartments. The inner mitochondrial membrane that invaginates to form crista and at the cristae junctions, the mitochondrial contact site and cristae organizing system (MICOS) complex resides (14,17,18). The MICOS complex bonds to the sorting and assembly machinery (SAM) complex (17). The MICOS complex is currently known to be made up of Mic10, Mic13, Mic19, Mic25, Mic26, Mic27, and Mic60 (17). Together, the interaction between the MICOS and SAM complexes connect the inner and outer mitochondrial membranes to form the mitochondrial intermembrane space bridging (MIB) complex (17). These proteins and their interactions are important for organizing and structuring the mitochondria to allow for maximum and efficient respiration and ATP generation (19). The MICOS complex, SAM complex, and OPA1 can all affect the dynamics of the cristae in varying ways (14,17,20,21). OPA1 is vital in promoting the formation of cristae and as it is responsible for determining the width of the cristae (14). Additionally, the MICOS subunits interact with the SAM complex to assemble β-barrel proteins and maintain contact between the outer and inner membranes (14,21,22). These all are necessary to maintain the overall shape and distances in mitochondrial networks. Overall, cristae membranes are crucial as they host the electron transport chain machinery and the ATP synthase dimers, which are necessary for the production of ATP through oxidative phosphorylation (9,20). However, given the critical relationship observed between mitochondrial morphology and function, determining the effects of SAM-associated gene knockout is necessary to understand the functional implications concomitant with changes in the mitochondrial ultrastructure and morphology.

While the functions of the SAM complex may easily be considered as a subset of the MIB complex, the SAM complex remains an interesting target distinct from the MICOS complex. The SAM complex is made up of three subunits, Sam35, Sam37, and Sam50 (23). While both Sam35 and Sam37 are peripheral membrane proteins that are not required for survival, the central component, Sam50, a 7–8 nm diameter β-barrel channel, interacts with the MICOS complex to modulate protein transport and mitochondrial morphology (17,21,22,24). Specifically, Sam50 stabilizes the MIB complex for protein transport, respiratory chain complex assembly, and cristae integrity regulation (25). In this way, it is understood that to structurally form and sustain the cristae, proteins of the MICOS complex assemble at the cristae junction and bind directly to Sam50 (19,21,26). Within the entire MIB complex, Sam50 is considered to be among the most important proteins, alongside members of the MICOS complex Mitofilin and Chchd3, for proper mitochondria structure and function (27). The importance of Sam50 is relevant, as loss of *Sam50* can result in diminished cristae count, reduced cristae dynamic events, abnormal cristae, and a lack of cristae junctions, the sites at which cristae are tethered to the mitochondrial inner boundary membrane (14). However, the implications of the loss of *Sam50* on mitochondrial dynamics in 3D is unknown.

2D microscopy techniques allow for the viewing of typical tubular mitochondria (28,29), however, 3D phenotypes of mitochondria differ between tissue types and disease states (10,30,31). Mitochondrial swelling can occur as a response to dysregulation of ion homeostasis (32), while toroid mitochondria can arise in a calcium-dependent manner in stress states (33). Through fusion and fission dynamics, it is understood that mitochondria can respond to cellular stress to balance factors including volume, which typically allows for greater ATP generation, and surface area, which typically allows for organelle-to-organelle contacts which are crucial for extraneous processes such as calcium homeostasis (10,34,35). Therefore, 3D microscopy techniques (34,36–38) are required for an understanding of many of the diverse phenotypes which mitochondria might display.

This study specifically examined the metabolomics of the knockdown of *Sam50* in murine and human myotubes. For the knockout of genes, myotubes, derived from satellite cell fusion, have emerged as robust cellular models which mimic the properties found in skeletal muscle (39). Importantly, human and murine myotubes remain distinct, with separate phenotypes and metabolic progression (40). Therefore, human myotubes were *ex vivo* isolated from skeletal muscle biopsies and differentiated in multinucleated myotubes in culture (41). To measure structure, we used a mixed-method approach utilizing principally serial block-face-scanning electron microscopy to allow for manual tracings to be performed on z-direction slices, known as orthos, which were then aligned in analysis software Amira (34), and reconstructed into 3D structures. Serial block-face-scanning electron microscopy (SBF-SEM) importantly allows for the ability to see mitochondrial complexity, as it offers high x- and y-resolution and range. We also sought to compare differences in mitochondrial structure and function based on human myotubes to better elucidate if *Sam50* may have a differential role in these models. Beyond structural dynamics, we sought to combine how changes in mitochondrial morphology upon *Sam50*-deficinecy alters mitochondrial function and general biochemical pathways, which may offer further implications for therapeutical targets involved in *Sam50*.

## Materials and Methods

### Mice Care

All procedures for the care of mice were in accordance with humane and ethical protocols approved by the University of Iowa Animal Care and Use Committee (IACUC) which follows the National Institute of Health (NIH) Guide for the Care and Use of Laboratory Animals, as described previously (42). Experiments utilized WT male C57Bl/6J mice housed at 22 °C on a 12-hour light/12-hour dark cycle with free access to water and standard chow. Mice were anesthetized with 5% isoflurane/95% oxygen.

### Murine and Human Myotube Isolation

Satellite cell isolation was performed as previously described (42). Satellite cells from C57Bl/6J mice were plated on BD Matrigel-coated dishes and activated to differentiate into myoblasts and myotube. Specifically, for myoblast differentiation, mouse-derived satellite cells with Dulbecco’s modified Eagle medium (DMEM)-F12 containing 20% fetal bovine serum (FBS), 40 ng/ml basic fibroblast growth factor, 1× non-essential amino acids, 0.14 mM β-mercaptoethanol, 1× penicillin/streptomycin, and Fungizone. then maintained with 10 ng/ml basic fibroblast growth factor. When myoblasts were 90% confluent, they were differentiated to myotubes in DMEM-F12 containing 2% FBS and 1× insulin–transferrin–selenium. Three days after differentiation, myotubes were infected with 1 µg CRISPR/Cas9 (Santa Cruz Sam50 CRISPR Plasmid; sc-427129) to achieve Sam50 deletion, which was validated by qPCR. Experiments were performed 3-7 days after infection.

### RNA Extraction and RT-qPCR

The RNA was isolated with an RNeasy kit (Qiagen Inc) and quantified by measuring absorbance at 260 nm and 280 nm with a NanoDrop 1000 spectrophotometer (NanoDrop products, Wilmington, DE, USA). Isolated RNA (∼1µg) was reverse transcribed with a High Capacity cDNA Reverse Transcription Kit (Applied Biosciences, Carlsbad CA) and amplified by real-time quantitative PCR (qPCR) with SYBR Green (Life Technologies, Carlsbad, CA) (43). Three for each qPCR, t samples (∼50 ng DNA each) were placed in a 384-well plate and underwent thermal cycling in an ABI Prism 7900HT instrument (Applied Biosystems). Thermal cycling conditions were set as follows: 1 cycle at 95°C for 10 min; 40 cycles of 95°C for 15 s, 59°C for 15 s, 72°C for 30 s, and 78°C for 10 s; 1 cycle of 95°C for 15 s; 1 cycle of 60°C for 15 s; and one cycle of 95°C for 15 s. Results were normalized to glyceraldehyde-3-phosphate dehydrogenase (*GAPDH*) and presented as relative fold changes.

### Measurement of OCR Using Seahorse

Oxygen consumption rate was measured for *Sam50* KD fibroblasts using an XF24 bioanalyzer (Seahorse Bioscience: North Billerica, MA, USA) as previously described (42,44). Briefly, cells were plated at a density of 20 × 10^3^ per well and differentiated for 3 days. For *Sam50* KO models, three days after differentiation, myotubes were transfected with CRISPR/Cas9 to achieve KO as described above. Cells were incubated in XFLDMEM (supplemented with 1 g/L D-Glucose, 0.11 g/L sodium pyruvate, and 4 mM L-Glutamine), without CO_2_ for 60 minutes. Cells were subsequently treated with mitochondrial stress modulators oligomycin (1 μg/ml), carbonyl cyanide 4-(trifluoromethoxy)phenylhydrazone (FCCP; 1 μM), rotenone (1 μM), and antimycin A (10 μM), in that order, while remaining in the XF-DMEM media and assessed using an XF24 Seahorse Analyzer. After analysis, cells were lysed in 20μl lysis buffer (10mM Tris, 0.1% TX-100, pH 7.4) (44),and 480 μl of Bradford reagent. was added to each well as described previously (40). Total protein concentration was measured at an absorbance at 595 nm and used for normalization. For each sample, three independent experiments were performed for each condition with representative data from the replicates being shown.

### Segmentation and Quantification of 3D SBF-SEM Images Using Amira

SBF-SEM ortho slices were 3D reconstructed using contour tracing (manual segmentation) in Amira to perform 3D reconstruction, as described previously (34,45). 300−400 slices ortho slices were transferred to Amira, and 50-100 serial sections were chosen, stacked, aligned, and visualized. A blinded individual familiar with organelle morphology traced structural features manually on sequential slices of micrograph blocks. Dots represent the total number of mitochondria surveyed, which varies.

### Gas Chromatography-Mass Spectrometry (GC-MS) and Analyzing Metabolomic Data

Samples were extracted for metabolites and prepared as previously designed (46) and performed at University of Iowa High Resolution Mass Spectrometry Facility according to standard procedures. TraceFinder 4.1 was used preloaded with standard verified peaks and retention times to compare metabolite peaks in each sample against an in-house library of standards. to correct for drift over time by using QC samples, we used previously described protocols (47). All data were normalized to an internal standard to control for extraction, derivatization, and/or loading effects. Metabolomic analysis was performed as described previously (46) using the web service MetaboAnalyst 5.0 (https://www.metaboanalyst.ca/MetaboAnalyst/ModuleView.xhtml, last accessed on 1 September 2022). All tests performed were using built-in comparison, including one-way ANOVA and Fisher’s LSD multiple comparison test. The fold enrichment number was calculated by the observed hits by the expected hits, which were determined by MetaboAnalyst 5.0.

### Data Analysis

In graphs, black bars represent the standard error, and dots represent individual data points. An unpaired t-test was used for data with only two groups while ANOVA with Tukeys’ *post-hoc* analysis was used for data with two or more groups. For both types of analyses, GraphPad Prism software package was used (La Jolla, CA, USA). A minimum threshold of p < 0.05 indicated a significant difference. Higher degrees of statistical significance (**, ***, ****) were defined as p < 0.01, p < 0.001, and p < 0.0001, respectively.

## Results

### Loss of Sam50 causes Loss of Mitochondrial Size and Alterations in Morphology in Murine and Human Myotubes

Given that mitochondrial size is linked with respiratory efficiency (31), we first sought to determine if mitochondrial size changed as a result of *Sam50* ablation. Following CRISPR/Cas9 knockout of Sam50 (Figure 1A), SBF-SEM micrographs were obtained (Figure 1B), and we performed manual contour segmentation for 3D reconstruction (Figure 1C-D) and subsequent quantification (Figure 1E). For each experimental condition, 10 myotubes were used with sectioning performed on 10 µm by 10 µm slices with a z-directional depth of 300 µm (Figure 1B). Fifty ortho slices (Figure 2A-B) were surveyed and manual contour tracing was performed on each mitochondrion to perform 3D reconstructions (Figure 2A’-B’). This allowed for observation of the complete mitochondrial 3D structure (Figure 2A’’-B’’) in murine myotubes as well as human myotubes (Figure 2A’’’-B’’’’’) (48). In each experimental condition, 3 regions of interests were considered, within which approximately 250 mitochondria were quantified for a total of 750 mitochondria (Figure S1).

**Figure 1:**
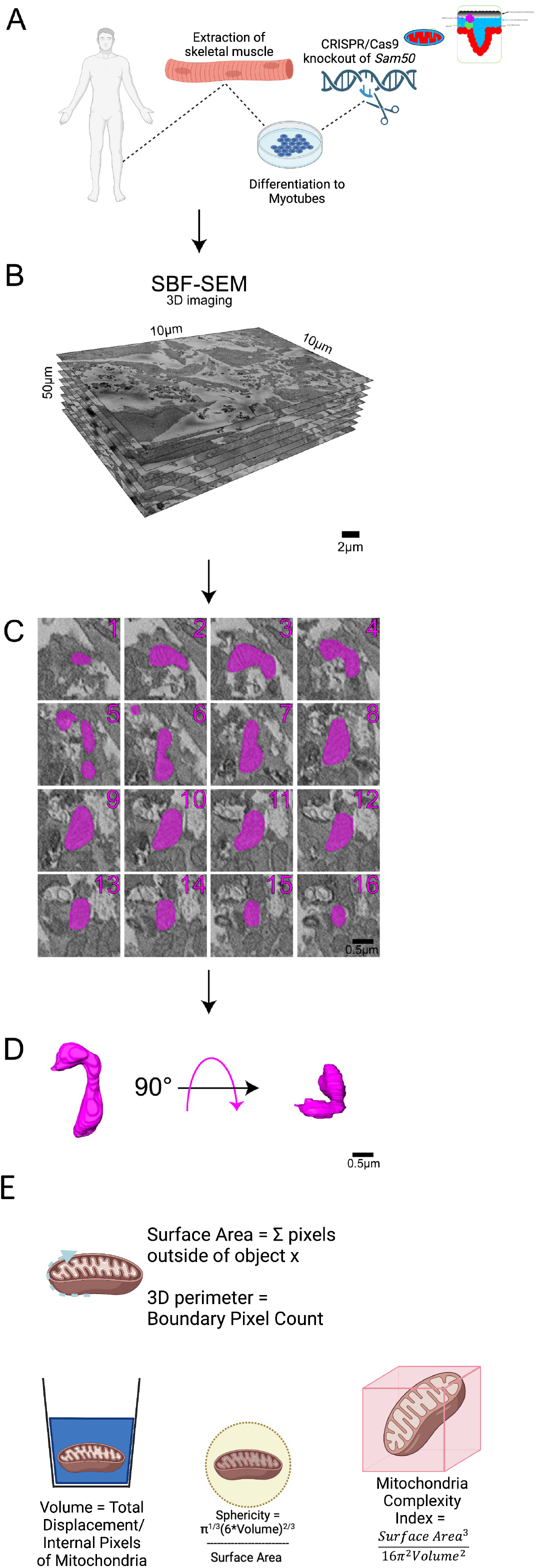
Workflow of SBF-SEM manual contour reconstruction to create 3D mitochondrial structure upon loss of *Sam50*. (A) Workflow depicting isolation of *Sam50*-deficient human myotubes. SBF-SEM allows for (B) ortho slice alignment. (C) Manual segmentation of ortho slices was performed to yield (D) 3D reconstructions of mitochondria. (E) Once 3D reconstruction was performed, quantifications were performed for mitochondria.

**Figure 2:**
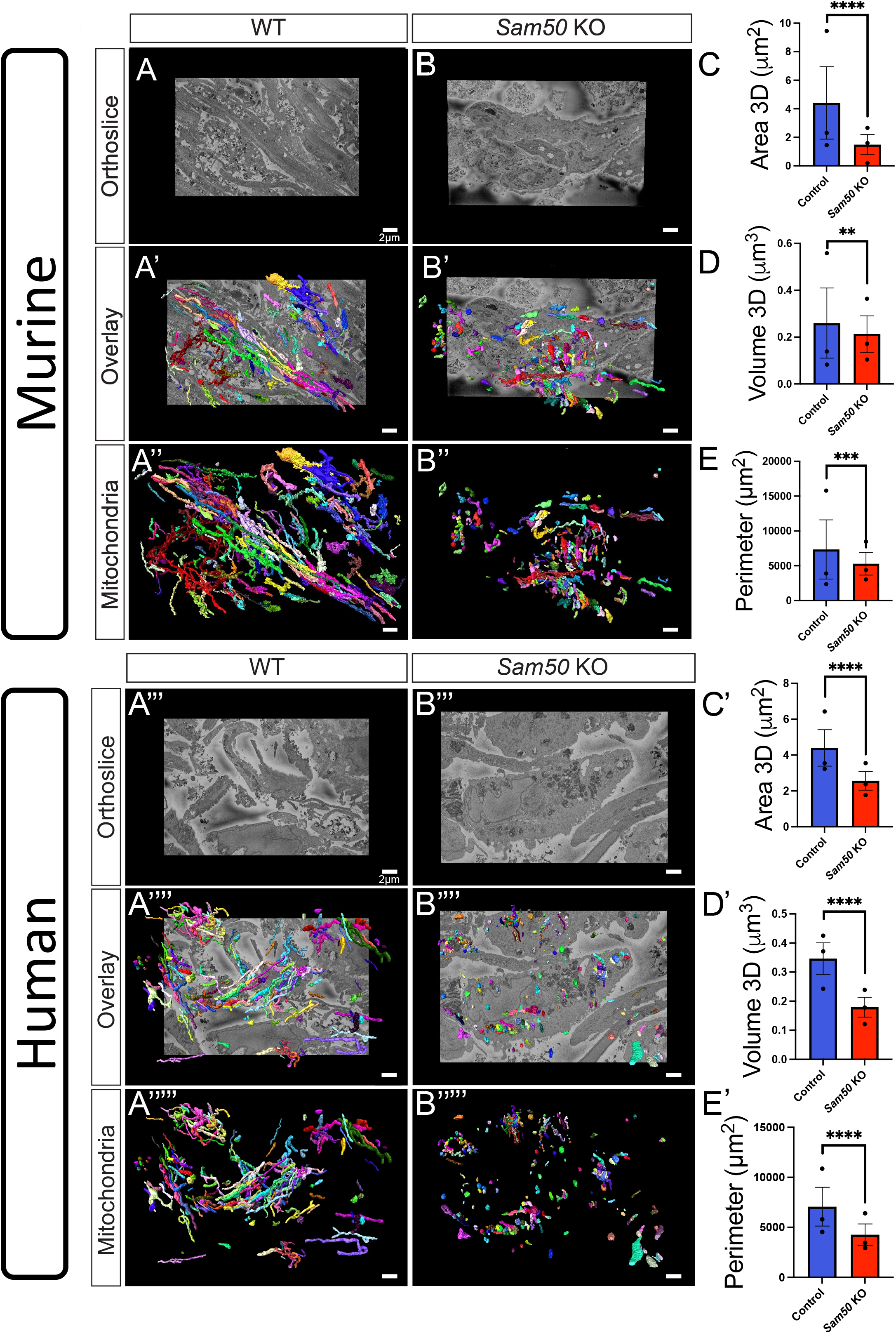
SBF-SEM shows changes in human and murine myotubes mitochondrial 3D structure upon loss of *Sam50.* (A) Control and (B) *Sam50* knockout murine myotubes orthoslice, displayed. Approximately 50 of these orthoslices are segmented to perform 3D reconstruction. (A’) Control and (B’) *Sam50*-deficient myotubes 3D renderings displayed overlaid on SBF-SEM orthoslice. (A’’) Isolated 3D mitochondria from control and (B’’) *Sam50*-deficient myotubes. (C) Upon loss of Sam50, on average mitochondria have reduced volume, (D) surface area, (E) and perimeter. (A’’’) Control and (B’’’) *Sam50* knockout human myotubes orthoslice, displayed. Approximately 50 of these orthoslices are segmented to perform 3D reconstruction. (A’’’’) Control and (B’’’’) *Sam50*-deficient myotubes 3D renderings displayed overlaid on SBF-SEM orthoslice. (A’’’’’) Isolated 3D mitochondria from control and (B’’’’’) *Sam50*-deficient myotubes. (C’) Upon loss of *Sam50* in human myotubes, on average mitochondria have reduced volume, (D’) surface area, (E’) and perimeter.

We observed that mitochondrial fragmentation was increased in *Sam50*-deficient murine myotubes compared to control myotubes (Figure 2C). Similarly, the area in 3D was also decreased upon *Sam50* loss (Figure 2D). This mimicked a loss in perimeter (Figure 2E), reducing the total mitochondrion surface area. In considering *Sam50*-deficient human myotubes (Figure 2C’-E’), we noticed a similar phenotype with loss of mitochondrial volume, area, and perimeter when compared to human myotubes (Figure 2C’-E’). From there, we sought to understand how beyond only size, mitochondria may differentially change in shape in murine and human myotubes following loss of *Sam50*.

To begin with, we showed mitochondria, from the transverse plane, in control (Figure 3A-A’) and *Sam50-*deficient murine myotubes (Figure 3B-B’), as well as human myotubes (Figure 3A’’-B’’’). Viewing mitochondria from both of these planes showed that at baseline murine myotubes are much more complex than human myotubes. While both murine and human myotubes showed loss in mitochondrial structure following loss of *Sam50*, this was a more significant loss in murine myotubes. Next, we sought to determine whether mitochondrial sphericity was altered. *Sam50*-deficiency was associated with an increase in mitochondrial sphericity which is indicative of reduced complexity in murine (Figure 3C) and human myotubes (Figure 3C’). To further understand mitochondrial complexity, we measured the mitochondrial complexity index which is a measure of the surface area-to-volume ratio. Mitochondria become significantly less complex upon loss of *Sam50* (35) in murine (Figure 3D) and human myotubes (Figure 3D’). This was further validated with mito-otyping, an approach that involves organizing mitochondria by volume in a similar manner to the arrangement of chromosomes during karyotyping. With mito-otyping, we observed that while control mitochondria were highly elongated *Sam50-*deficient mitochondria generally become less elongated, resulting in a decrease in overall volume in both murine (Figure 3F) and human (Figure 3F’) myotubes. Overall, loss of *Sam50* has been shown to be associated with a dysregulation of mitochondrial structure. Considering the dramatic decrease in mitochondrial size observed in *Sam50*-deficient myotubes, we sought to also investigate whether these morphological effects influenced the efficiency of mitochondrial functions.

**Figure 3:**
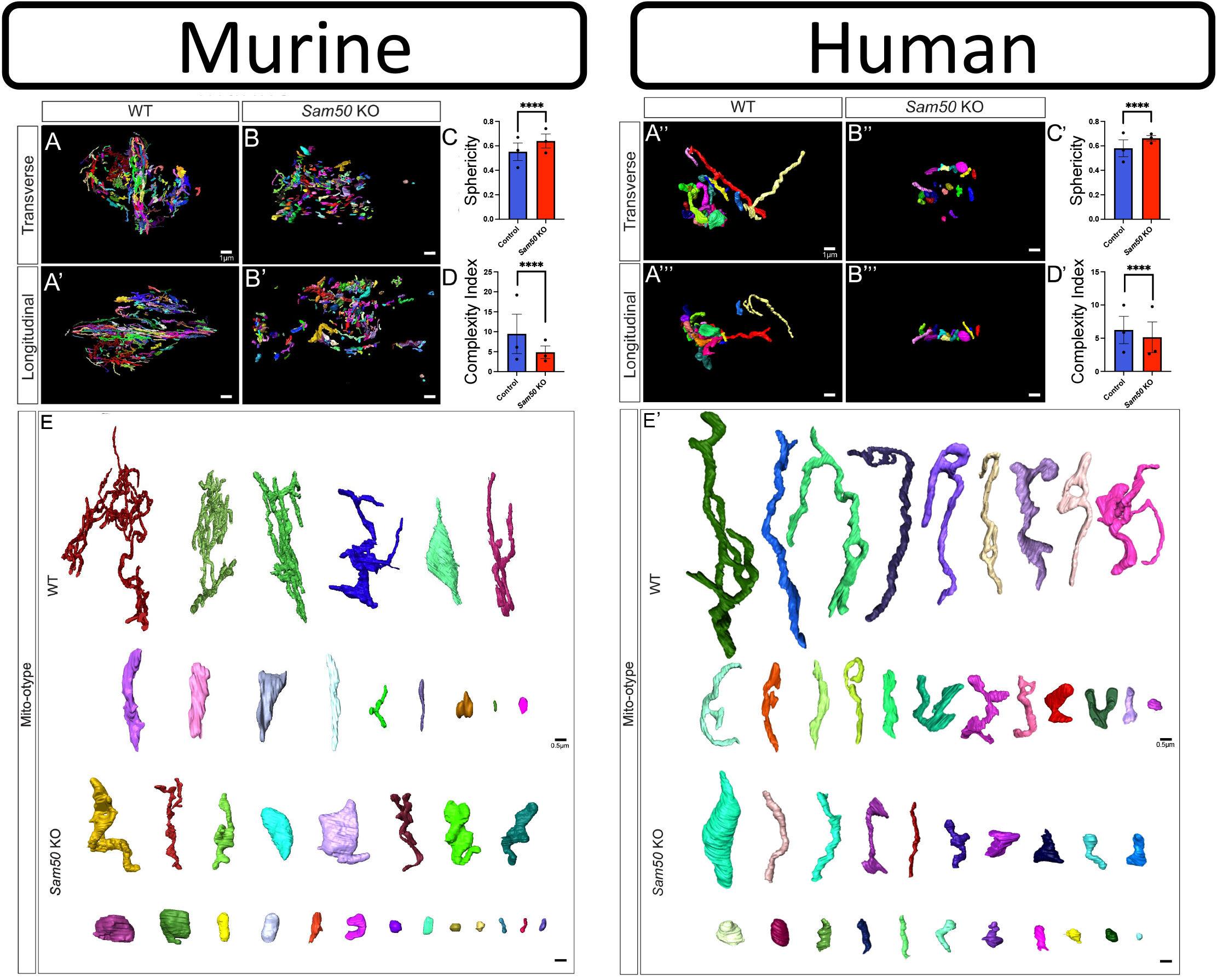
Changes in human and murine myotubes mitochondrial branching and networking upon loss of Sam50 revealed in SBF-SEM. (A) 3D Reconstructions showing control and (B) *Sam50*-deficient murine and human (A’’-B’’) myotubes from a transverse-point of view. Murine (A’) and human (A’’’) 3D Reconstructions showing control and (B’) *Sam50*-deficient murine and human (B’’’) myotubes from a longitudinal point of view. (C) Sphericity of mitochondria in control and *Sam50*-deficient murine and (C’) murine myotubes. (D) Mitochondrial complexity index (MCI), which is analogous to sphericity, comparing control and *Sam50*-deficient murine and (D’) human myotubes. (E) Mito-otyping, to display the diversity of mitochondrial phenotypes as ordered by volume, in control and *Sam50*-deficient murine and (E’) human myotubes.

### Increased Autophagosomes Occurs Upon Loss of *Sam50* in Human Myotubes

Most abundantly, macroautophagy is the routine cellular mechanism for organelles, including mitochondria, through principally autophagosomes (49). Targeting of mitochondria in autophagy, known as mitophagy, can serve as a critical cellular response to reactive oxygen species or other forms of dysfunctional mitochondria; as such, mitophagy has been proposed as a key target in limiting the pathological progression of neurodegenerative diseases (49). *Sam50* may play a crucial role in the biochemical triggering of these mitophagy events. Of note, *Sam50* interacts with autophagy receptor p62 (50). Beyond this, past research has also found that *Sam50* deficiency results in autophagic flux, with mtDNA protecting from complete mitochondrial degradation (51). However, autophagy remains a dynamic process that may be more effectively studied using 3D reconstruction (52), and the effects of *Sam50*-deficiency on the structure of autophagic organelle machinery are poorly elucidated.

Since humans and murine myotubes reflected mitochondrial structural arrangements with loss of *Sam50*, we focused on human myotubes. Here, again using 3D reconstruction from 10 µm by 10 µm slices obtained from 10 human myotubes, with a z-directional depth of 300 µm, we surveyed 50 ortho slices (Figure 4A). Following manual segmentation of these 50 ortho slices, 3D renderings of autophagosomes were able to be created for control and *Sam50*-deficient human myotubes (Figure 4B-B’). We found that autophagosomes, when normalized to the same area regions of interests (ROI), were much higher in count upon loss of *Sam50* in human myotubes (Figure 4C). Beyond only count, the average volume of each autophagosome increases (Figure 4D). These increases in overall size can be illustrated when autophagosomes are organized on the basis of their volume (Figure 4E). This suggests larger autophagosomes that may be more active or have increased cargo loads. Together, these results show that autophagosome formation was increased in *Sam50*-deficient myotubes compared to control myotubes, suggesting an uptick in mitophagy.

**Figure 4:**
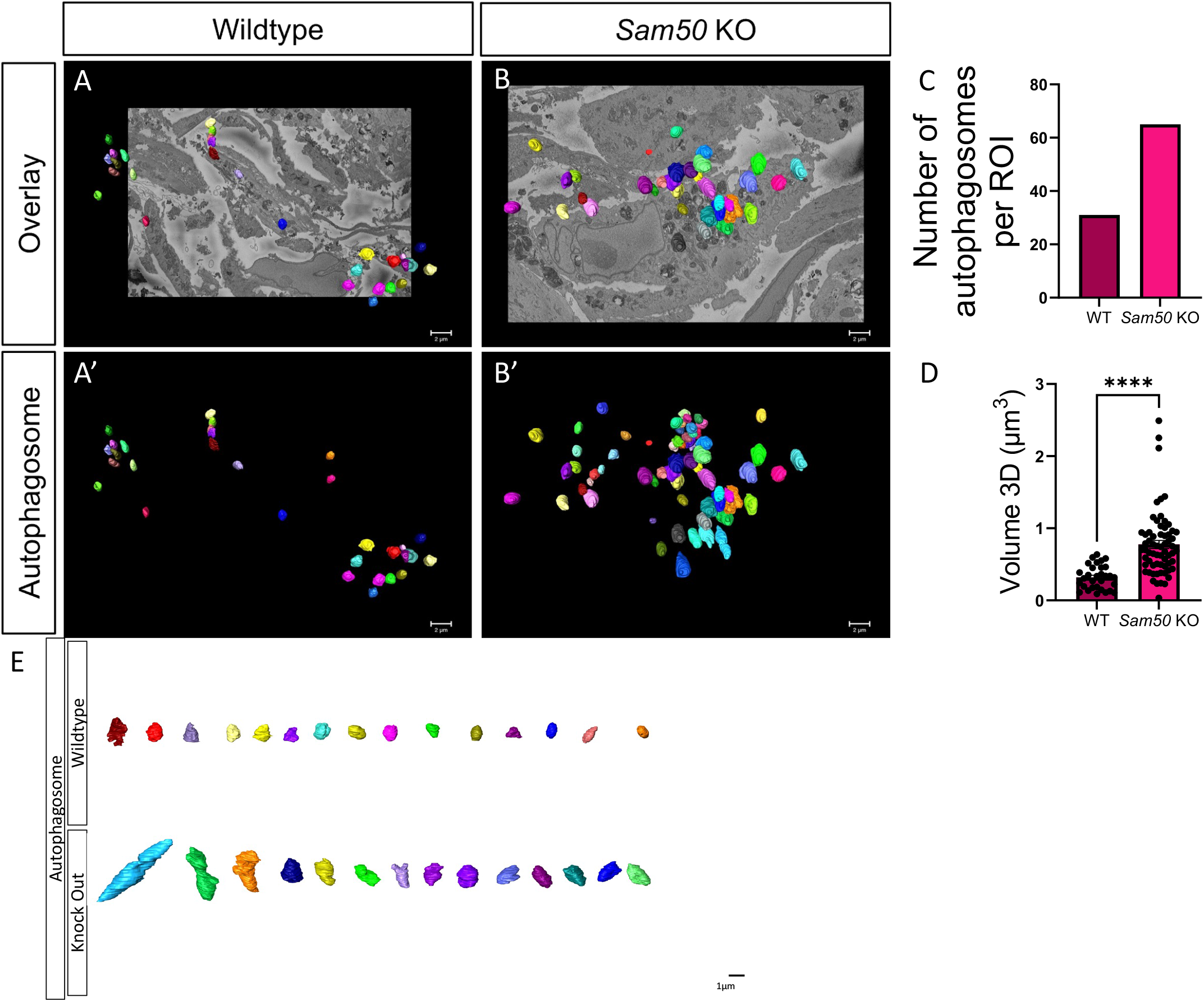
Changes in macroautophagic autophagosomes count and volume upon loss of *Sam50* revealed in human myotubes through SBF-SEM. (A) 3D Reconstruction overlaid on z-directional orthoslice of autophagosomes in control myotubes and (B) *Sam50* knockout (KO) myotubes. (A’) Isolated 3D reconstruction of autophagosomes in control and (B’) Sam50 KO myotubes. Quantifications of (C) the number of autophagosomes and (D) the average volume of autophagosomes upon loss of *Sam50* in human myotube. (E) Autophagosomes organized on the basis of their volume going from left to right to illustrate the diversity of recycling machinery.

### Alterations of Mitochondrial Efficiency Occurs in *Sam50*-deficient Murine and Human Myotubes

To investigate whether observed structural changes mitochondria influenced mitochondrial efficiency, we assessed the overall oxidative capacity of control and *Sam50-* deficient human and murine myotubes. For these experiments, we used an XF24 Seahorse Analyzer which measures oxygen consumption rate (OCR), an indicator of mitochondrial oxidative photophosphorylation, across various drug applications. Prior to mitochondrial efficiency studies, knockout of Sam50 was confirmed in both murine and human myotubes (Figure 5A-4A’). Overall, *Sam50*-deficient murine and human myotubes showed reduced OCR at all time intervals (Figure 4B-B’). To determine basal OCR, cells were assessed under normal oxidative conditions. Basal OCR was impaired in cells lacking Sam50, compared to control cells (Figure 5C-C’). A series of oxidation modulators were applied to further investigate the effects of *Sam50* on oxidative processes. First, oligomycin was applied to inhibit ATPase and induce hyperpolarization of the membrane, which allowed us to examine the role of *Sam50* in ATP-linked respiration (Figure 5D-D’). Carbonyl cyanide-4 (trifluoromethoxy) phenylhydrazone (FCCP), a mitochondrial oxidative phosphorylation uncoupling agent, was then applied. With FCCP treatment, we show that under conditions of maximal respiration, OCR is reduced with the loss of *Sam50* (Figure 5E-E’). compared to control Finally, myotubes were treated antimycin-a and rotenone inhibitors electron transport, which allowed for the glycolytic reserve to be determined as a function between maximum OCR and baseline OCR (Figure 5F-F’). Overall, with our mitochondrial functional assay, we demonstrate that oxidative capacity is impaired upon ablation of *Sam50* in murine and human myotubes.

**Figure 5:**
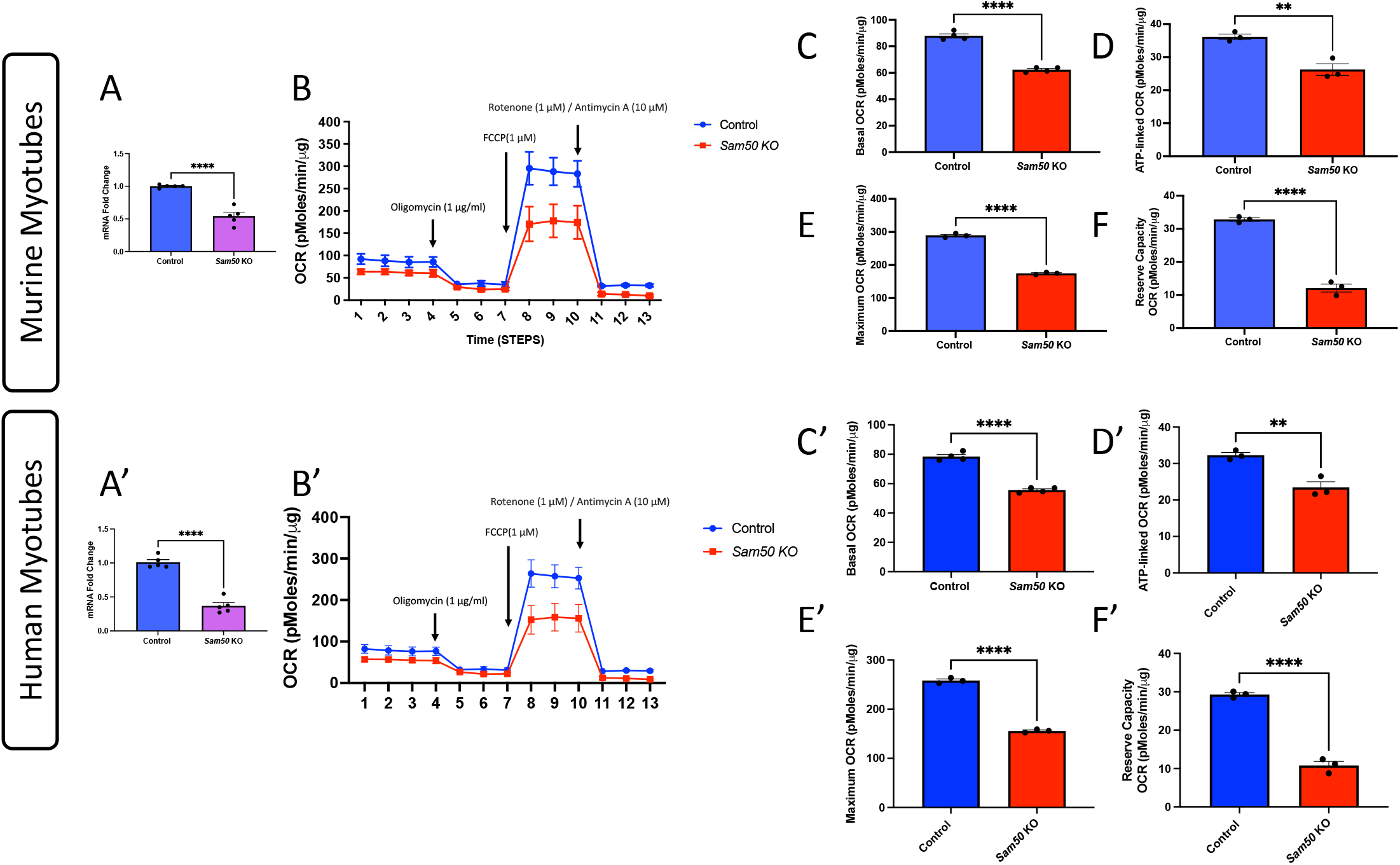
XF24 Seahorse Analyzer measured respiration rate in murine and human myotubes. Validation through quantitative polymerase chain reaction of knockout of *Sam50*. (A) Decreased mRNA transcripts of *Sam50* in murine myotubes and (A’) human myotubes following CRISPR/Cas9 transient knockout. (B) *Sam50*-deficient murine and (B’) human myotubes showing reduced oxygen consumption rate (OCR) in comparison to control. (C) The first 4 time points measure basal respiration, or mitochondrial OCR at baseline in murine and (C’) human myotubes. (D) The next three time points following application of oligomycin (1 µg/mL) shows decoupling of respiration dependent on ATP in murine and (D’) human myotubes. (E) The next three time points measured following the addition of carbonyl cyanide 4-(trifluoromethoxy)phenylhydrazone (FCCP; 1 μM) shows OCR under maximum respiration in murine and (E’) human myotubes. (F) The final three time points show OCR following rotenone (1 μM) and antimycin A (10 μM) show minimum OCR that is not dependent on mitochondria in murine and (F’) human myotubes.

### *Sam50*-deficiency in Human Myotubes Changes Metabolomic Pathways

Since human and murine myotubes had similar structural changes with the loss of *Sam50*, we specifically looked at human myotubes to investigate the specific biochemical changes resulting from the loss of *Sam50*. To investigate the metabolic effects of *Sam50* loss, we employed Gas Chromatography-Mass Spectrometry-based (GC-MS) metabolomics. Analysis using a volcano plot revealed significant downregulation of ß-alanine, gamma aminobutyric acid (GABA), and hypotaurine upon *Sam50* loss (Figure 6A). Heat map analysis further showed downregulation of various metabolites, including amino acids such as glutamine, phenylalanine, glutamate, threonine, and proline, suggesting a decrease in amino acid products due to increased amino acid metabolism. On the other hand, ribose, ß-Hydroxy, indolepropionic, inotisol, and dihydrophenylalanine were upregulated (Figure 6B). Enrichment analysis confirmed these findings, with significant enrichment observed in the metabolism and degradation of amino acids and fatty acids. Additionally, the metabolomic analysis revealed an increase in amino acid metabolism and fatty acid metabolism (Figure 6C). Collectively, these findings suggest that altered biochemical pathways may arise in response to, or as a mechanism of, the disrupted 3D structure of mitochondria upon *Sam50* loss.

**Figure 6:**
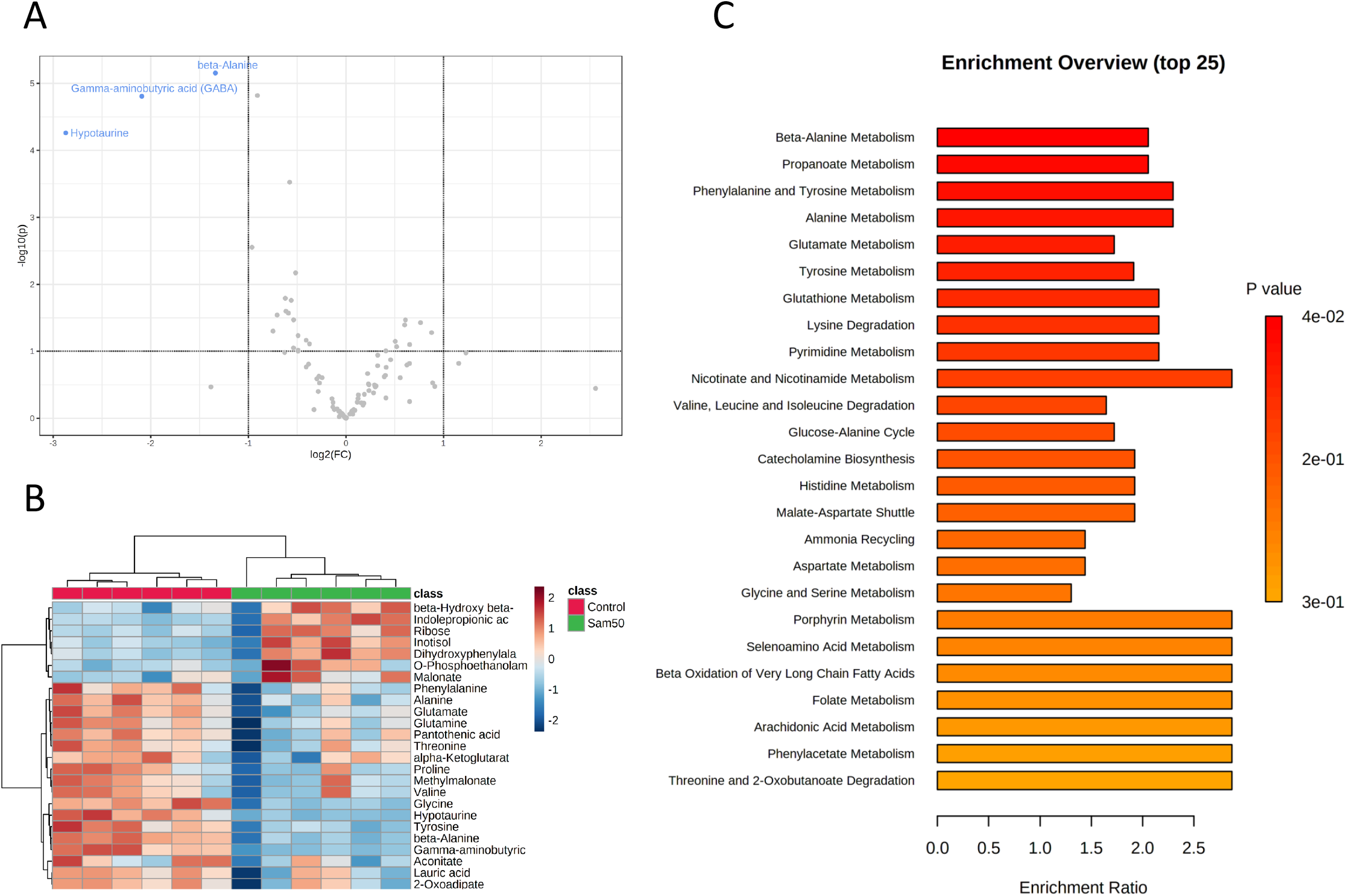
Metabolomic Comparison of *Sam50*-deficient human myotubes and control myotubes. (A) Volcano plot of metabolites that were differentially regulated in *Sam50*-deficient myotubes compared to WT samples. (B) Relative abundance of ions in *Sam50*-deficient versus control myotubes visualized on a heatmap. (C) Enrichment analysis displaying enriched metabolites in Sam50-*deficient* myotubes.

## Discussion

Together, these data suggest Sam50 is critical for the establishment and maintenance of mitochondria, mitochondrial cristae structure, and mitochondrial metabolism. To our knowledge, we are the first group to look at the 3D structure of mitochondria in murine and human myotubes following the genetic knockdown of *Sam50*. Previously, transmission electron microscopy has been utilized to observe that Sam50 can be used as a therapeutic target, with overexpression rescuing mitochondrial morphology upon injury-induced loss of morphology (53). Interestingly, our results show that *Sam50* may be functionally required for many of the diverse phenotypes previously observed in other tissue types (10). Loss of *Sam50*, importantly, showed a loss of mitochondrial morphology diversity, which may have functional impacts on functions including calcium homeostasis and mitochondria-organelle contacts.

Previously, studies have shown that Sam50 levels may rise in response to mitochondrial dysfunction, suggesting it acts as a sort of recovery molecule (53). This highlights the importance of understanding how Sam50 interacts with the MICOS complex to influence overall mitochondrial structure. While it is well understood that the MICOS structure modulates cristae changes, here we also show a structure loss in mitochondria structure upon loss of the SAM complex. It is possible this occurs due to the interaction of the MICOS complex, which may also affect overall mitochondrial dynamics, potentially through the modulation of fusion and fission dynamics such as DRP1 and OPA1, which also affects cristae structure (13). Future experiments may look at cristae to understand how cristae structural rearrangement alters 3D mitochondrial general phenotype in the loss of the MIB complex. Therefore, future experiments should consider further utilizing focused ion beam-scanning electron microscopy, which allows for machine-learning outputs of 3D cristae morphology to be constructed (34,36,54).

In this study, we use 3D microscopy to dissect changes in mitochondrial morphology and function in myotubes with altered *Sam50* expression. From images captured with this technique, we demonstrated mitochondrial volume concomitantly occurs with a loss of complexity in both murine and human myotubes. By studying both human and murine myotubes we were able to *show* how murine myotubes mitochondria are relatively larger than human myotubes, but loss of Sam50 cause mitochondria to undergo significant reductions in size and morphology in both models. It has been suggested microscopic imaging can be targeted for the development of treatments to correct cellular defects (53). Specifically, overexpression of Sam50 was shown to rescue ischemia/reperfusion injury-induced changes by Yin and colleagues in a neural model (53), highlighting Sam50 may be an important therapeutic in a mitochondrial-dependent manner.

Importantly, our results show the importance of understanding the functional impact of mitochondria decreasing in elongation, as past mechanistic studies remain conflicting. Previous studies have shown that mitochondrial elongation may be indicative of reduced fission dynamics, such as due to the loss of Drp1 (55). Mitochondria dysfunction may also arise as a result of mtDNA mutations (7). Interestingly, in a Drp1-dependent manner, mtDNA mutations may cause mitochondrial morphological alterations in Parkinson’s disease; however, knockdown of Drp1 can rescue mitochondria through elongation (55). Elongated mitochondria can also serve a distinct function to distribute membrane potential across a wider area (10). Past studies have shown that loss of the mitochondrial membrane potential alters mitochondria volume (56). Membrane potential is also regulated in cristae dynamics (57), suggesting alteration in the MIB complex may affect membrane potential. However, past research has shown that *Sam50* downregulation does not drastically alter membrane potential (25). Beyond this, it has also been suggested that elongated mitochondria offer increased ATP production and protect against mitophagy, which may aid in explaining the uptick in autophagy observed following *Sam50* KO (58).

Outside of elongated mitochondria, a key mitochondrial characteristic observed exclusively in wild-type myotubes is a diversity of phenotypes. Interestingly, there is greater complexity in murine myotubes but reduced elongation when compared to murine myotubes. For both myotubes, following *Sam50* KO, many phenotypes have much higher complexity (Figure 3D), as well as higher surface area. This is epitomized by toroid-shaped mitochondria, which offer mitochondria an increased surface area, at the expense of volume for cristae (10). Although calcium-dependent donut mitochondria have been observed in states of ROS stress and disease (10,33), they may also serve beneficial tissue-dependent roles (59). Their absence in *Sam50-deficient* models may signify a potential inhibition of calcium. Beyond this, their increased surface area may allow wild-type myotubes to have increased contact sites with the endoplasmic reticulum, lipid droplets, and other organelles. This remains relevant as mitochondria endoplasmic reticulum sites (MERCs) serve a multitude of functions including calcium homeostasis (60,61) and regulating autophagy as autophagosomes form at ER sites (52,62). Future studies may consider further exploring this link by understanding how mitochondrial contact sites may change upon loss of *Sam50*.

Also offering valuable clues about what may be causing mitochondria function to be affected is metabolic changes we observed using GC-MS. Many of the results from the metabolic analysis show that loss of *Sam50* may result in metabolic changes that parallel disease states. For example, elevations in levels of 3,4-dihydroxyphenylalanine, which we observed increased in *Sam50*-deficient myotubes (Figure 5B), are understood to occur in association with oxidative stress during disease states (63,64). Past results have also shown that *Sam50*-deficiency is associated with liver disease through increased lipid accumulation due to increased fatty acid oxidation (65). We noted a similar effect as beta-oxidation of fatty acids increasingly occurred in *Sam50*-deficiency. In the human heart, energetic demands are often carried out through the beta-oxidation of fatty acids (66). Therefore, these results suggest that it is possible that fatty acid oxidation is increased upon loss of *Sam50* to allow for energy production in states when mitochondria lose optimal functioning; however, future studies may further explore if in mitochondrial disease states, fatty acid oxidation changes in a *Sam50*-dependent manner.

Of note, we saw largely significant reductions in levels of GABA, hypotaurine, and ß-alanine in *Sam50*-deficient myotubes (Figure 5A), which may have several potential implications. Beyond functioning in embryonic neurogenesis and as a neurotransmitter, GABA acts in a potential mTOR-dependent manner with accumulation resulting in mitophagy and abnormal morphology (67). Typically GABA is metabolized by a pathway known as the GABA shunt, found in the mitochondrial matrix; decreased frequency of GABA may be linked to upregulated upstream targets in the GABA shunt (68). It is well understood that the primary role of hypotaurine is as a precursor for taurine when reacted with hydrogen peroxide (69). Critically, taurine serves numerous functions beyond antioxidant roles, protecting against mitochondrial disease, potentially through posttranslational modifications of mitochondrial tRNAs (70,71). Together, these studies suggest a potential antioxidant role of hypotaurine in mitochondria (72). Similar to hypotaurine, ß-alanine is also a precursor to taurine production. Studies have suggested that ß-alanine supplement increases skeletal muscle carnosine content, which affects calcium homeostasis and offers antioxidant abilities, which can increase muscle endurance, yet these results remain debated (73,74). Interestingly, however, increased ß-alanine has also been shown to result in mitochondrial fragmentation and oxidative stress (75). Overall, results across prior studies show varied results suggesting a tissue-dependent role of ß-alanine which requires further elucidation.

These reductions in ß-alanine parallel a larger trend of amino acid metabolism, evidenced by decreased metabolites of many amino acids (Figure 5B). This can have numerous implications on the dependency of *Sam50* for overall mitochondrial health during disease states. For example, the catabolism of fatty acids and amino acids is commonly observed as a method to bolster cancer growth (76). These changes may also be affecting the mTORC pathway (77). Recently it was found that the mTORC pathway interacts in an OPA-1 mediated manner and may be associated with the overall mitochondrial structure in stress states (78). Indeed, it has been found that the ATF4-mediated integrated stress response (ISR) is responsible for mediating cellular responses to oxidative stress and regulating amino acid metabolism (79). This pathway may be activated by ER or mitochondrial stress-dependent eIF2α phosphorylation (80). Future studies may aim to understand if metabolite pathways are changing in response to the structural-related declines in respiratory efficiency, or if the inverse is happening.

Alongside metabolic changes, mitochondrial structure may be related to alterations in autophagy that we observed. Upregulation of autophagy may be targeting ROS in mitochondria, which is often the cause of mitophagy activation (81,82). Interestingly, past research has found that cytotoxic ROS-dependent molecules enter the mitochondria in a Sam50-mediated manner to cause ROS build-up or apoptosis (83). Therefore, loss of Sam50 may serve a cytoprotective effect by preventing apoptosis and increasing proliferation in cancer cells (84). Beyond this, *Sam50* loss also contributes to an excess of ROS species (24,84), which contributes to many pathologies. Therefore, restoring *Sam50* may act as a mechanism to prevent ROS overproduction (84) and restore mitochondrial and cristae structure (53).

Together, our results additionally show that *Sam50* loss may further contribute to pathology through impairing mitochondrial structures, such as toroid shape and elongated mitochondria, reducing OCR, and altering metabolic pathways. Therefore, future research evaluating the targeting of *Sam50* to restore mitochondria 3D structure, function, and metabolism during *in vivo* disease states may prove a valuable future avenue.

## Contributions

B.S., M.K., A.O., and C.V. performed the experimental procedures, data analysis, interpretation of results, and writing of the manuscript. They share equal authorship.

F.Z., K.N., Z.V., and E.G.L. contributed to the experimental design and data analysis. A.K, Z.V., and E.G.L. J.Q.S., M.M., J.L., and Q.W. contributed to the data collection and analysis.

K.N., C.T.A., A.W., J.B., A.K., A.G.M., E.S., L.V., H.J.K., M.R.M., J.AG., S.D., B.C.J., A.T.C., S.M., D.S., and A.C. provided critical feedback and contributed to the writing and revision of the manuscript.

H.K.B and A.H. conceptualized the study, supervised the research, provided overall guidance, and revised the manuscript.

All authors reviewed and approved the final version of the manuscript.

## Data Sharing and Open Access

All data is available upon request to the corresponding author.

